# Molecular dissection of HERV-W dependent microglial- and astroglial cell polarization

**DOI:** 10.1101/2024.03.15.584408

**Authors:** Joel Gruchot, Laura Reiche, Luisa Werner, Felisa Herrero, Jessica Schira, Urs Meyer, Patrick Küry

**Affiliations:** Heinrich-Heine-University Düsseldorf, Medical Faculty and University Hospital Düsseldorf, Department of Neurology, D-40225 Düsseldorf, Germany; Institute of Veterinary Pharmacology and Toxicology, University of Zürich, Vetsuisse, Zürich, Switzerland; Neuroscience Center Zürich, University of Zürich and ETH Zürich, Zürich, Switzerland; Department of Neurology, Inselspital, Bern University Hospital and University of Bern, Bern, Switzerland

**Keywords:** Human endogenous retrovirus type W, neurodegeneration, neuroinflammation, multiple sclerosis, glia

## Abstract

The endogenous retrovirus type W (HERV-W) is a human-specific entity, which was initially discovered in multiple sclerosis (MS) patient derived cells. We initially found that the HERV-W envelope (ENV) protein negatively affects oligodendrogenesis and controls microglial cell polarization towards a myelinated axon associated and damaging phenotype. Such first functional assessments were conducted *ex vivo*, given the human-specific origin of HERV-W. Recent experimental evidence gathered on a novel transgenic mouse model, mimicking activation and expression of the HERV-W ENV protein, revealed that all glial cell types are impacted and that cellular fates, differentiation, and functions were changed. In order to identify HERV-W-specific signatures in glial cells, the current study analyzed the transcriptome of ENV protein stimulated microglial- and astroglial cells and compared the transcriptomic signatures to lipopolysaccharide (LPS) stimulated cells, owing to the fact that both ligands can activate toll-like receptor-4 (TLR-4). Additionally, a comparison between published disease associated glial signatures and the transcriptome of HERV-W ENV stimulated glial cells was conducted. We, therefore, provide here for the first time a detailed molecular description of specific HERV-W ENV evoked effects on those glial cell populations that are involved in smoldering neuroinflammatory processes relevant for progression of neurodegenerative diseases.

## 1. Introduction

Multiple sclerosis (MS) is a demyelinating disease of the central nervous system (CNS) leading to progressive neurodegeneration and irreversible cognitive and motor deficits. Despite extensive research, cause and pathomechanism of MS remain poorly understood. The classical concept of MS, described by early autoimmune mechanisms followed by a neurodegenerative phase, has repeatedly been challenged over the last years [1]. Meanwhile novel concepts predict the occurrence of smoldering neuroinflammation that leads to neurodegeneration already during early disease stages [2]. In this context and besides the influence of peripheral immune cells, this smoldering neuroinflammation was proposed to be primarily associated with reactive and neuroinflammatory microglia and astrocytes [3]. Advances in transcriptome analysis largely improved our understanding of expression pattern diversity and dynamics of these reactive- and disease-associated glial cells and multiomic approaches are currently used as state-of-the-art technology to further characterize these heterogeneous cell populations [4, 5].

Human endogenous retroviruses (HERVs) are evolutionary acquired DNA sequences and make up to 8% of the human genome, with most of them being epigenetically silenced or functionally irrelevant due to accumulation of mutations and/or truncations (summarized by [6]). Related to disease-initiating or -promoting pathological entities, Perron and colleagues reported in 1989 a viral entity isolated from leptomeningeal cell cultures of MS patients [7]. Later it was found that these viral particles belong to a human endogenous retrovirus of type W (HERV-W). Although anticipated as mainly inactive elements, it was proposed that certain environmental factors are able to activate these viral entities, leading to the expression of viral proteins. In this context, Herpesviridae, such as the Epstein Barr virus (EBV), have been identified to activate dormant HERVs [8, 9]. As EBV was recently revealed to be a leading cause of MS [10], this further corroborates a functional implication of HERV-W in the disease process. In MS, the envelope protein (ENV) of HERV-W was primarily found to be expressed and ENV protein levels, detected in patient blood and cerebrospinal fluid (CSF), were found to correlate with onset, progression, and severity of the disease [11-13]. Within the CNS, HERV-W ENV protein can be found as acellular deposits and to be expressed by myeloid cells whereas HERV-W ENV-positive astroglial- and lymphoid cells could also be detected [14, 15]. Functionally, it was proposed that HERV-W ENV exacerbates autoimmune responses [16], by activating peripheral immune- as well as endothelial cells [17-19] mediated via toll-like receptor-4 (TLR4)/Cluster of differentiation 14 (CD14)-signaling [19]. In 2013, a negative impact of the ENV protein on myelin repair activities by restricting the differentiation and maturation capacity of resident oligodendroglial precursor cells was described [20]. Moreover, we were recently able to show that HERV-W ENV activates microglial- and astroglial cells contributing to a neurotoxic environment featuring sustained demyelination, neurodegeneration and impaired remyelination [14, 21].

The strong impact of HERV-W on different CNS glial cell populations, particularly related to long-term damage processes relevant for MS progression, prompted us to investigate HERV-W-specific gene signatures of microglia and astroglia. These HERV-W ENV evoked responses were then compared to the transcriptomic patterns induced by lipopolysaccharide (LPS) stimulation, owing to the fact that both ligands can activate TLR-4.

## 2. Material and Methods

### 2.1. Primary rat glial cell culture

Isolation of primary rat microglial- and astroglial cells was performed conferring to the ARRIVE guidelines and in accordance with the National Institutes of Health guide for the care and use of Laboratory animals (NIH Publications No. 8023, revised 1978). The IRB (Institutional Review Board) of the ZETT (Zentrale Einrichtung für Tierforschung und wissenschaftliche Tierschutzaufgaben) at the Heinrich-Heine-University Düsseldorf approved all animal procedures under license O69/11. Both, microglial- and astroglial cells were isolated using magnetic activated cell sorting (MACS) from primary mixed glia cell cultures (as previously described in [14, 21]. Briefly, for the generation of mixed glial cell cultures, P0/P1 Wistar rat cortices were isolated, dispersed and cells were cultured on T-75 cell culture flasks in Dulbecco′s Modified Eagle′s Medium (DMEM; Thermo Fisher Scientific, Waltham, UK) substituted with 10% fetal calf serum (FCS; Capricorn Scientific, Palo Alto, CA, USA) and 2 mM L-glutamine (Invitrogen, Carlsbad, USA), 50 U/ml penicillin/streptomycin (Invitrogen, Carlsbad, USA). Medium change was performed every second day until cultures were grown confluently. After 10 days, flasks were transferred to an orbital shaker and shaken at 180 rpm/min at 37°C for 2h and then for another 22h. Microglia containing supernatants were collected at both time points, plated onto bacterial dishes and kept in the incubator (37°C, 5% CO2 and 90% humidity) allowing cells to attach. Remaining cells were dislodged using trypsin-EDTA (Gibco), spun down for 5 min at 300 × g and 4°C, supernatants were discarded and cell pellets were processed according to the manufacturers protocol for MACS of astroglial cells using the ACSA2 isolation kit (Miltenyi Biotec). Afterwards, 30.000 isolated astroglial cells per well were seeded in astrocyte medium (10% FCS, 2 mM L-glutamine, 50 U/ml penicillin/streptomycin in DMEM) onto 24-well plates. Three days post isolation, cells were stimulated with 1 µg/ml HERV-W ENV protein (Protein’eXpert, Grenoble, France) or 100 ng/ml lipopolysaccharide (LPS; Sigma-Aldrich, St. Louis, MO, USA) and respective buffer solutions (ENV buffer: Tris-HCL 20 mM, pH 7.5, 150 mM NaCl, 1.5% SDS, 10 mM DTT; LPS buffer: sterile H_2_O) serving as suitable controls for 24h.

For microglial purification a similar approach was used, in which microglial cells were detached from bacterial dishes using L-accutase (Gibco). The resulting cell suspension was spun down for 5 min at 300 × g and 4°C and processed for MACS using the CD11b/c microbeads according to the manufacturers protocol (Miltenyi Biotec). Subsequently, 300.000 microglia per well were seeded in microglia medium (10% FCS, 2 mM L-glutamine, 50 U/ml penicillin/streptomycin in DMEM) onto 24-well plates and stimulation experiments were performed 24 h after seeding using the same concentrations of HERV-W ENV protein and LPS, as used for astrocytes. Cell purities of microglial and astroglial cell cultures were analyzed regularly by Iba1/GFAP staining and were consistently at 98% for astrocytes and 99% for microglia. To avoid side effects through the recombinant production of HERV-W ENV protein, endotoxin levels were measured using the limulus amebocyte lysate-test and found to be below the detection limit (<5EU/ml) (as previously described in detail [14, 20, 21].

### 2.2. RNA preparation and bulk RNA Sequencing

For bulk RNA sequencing, stimulated microglial- and astroglial cells were lysed using β-mercaptoethanol supplied RLT buffer according to the manufacturer’s protocol (Qiagen, Hilden, Germany). Afterwards, total RNA was purified from cells using the RNeasy procedure (Qiagen, Hilden, Germany). Preparation of RNA library and transcriptome sequencing was conducted by Novogene Co. LTD (Bejing, China). Briefly, to generate RNAseq libraries, DNase digested total RNA samples were analyzed for quantity, integrity and purity using Agilent 5400 fragment analyser (Agilent Technologies, Inc. Santa Clara, USA). All samples in this study showed high quality RNA Quality Numbers (RQN; >9.3). Library preparation and bulk mRNA sequencing was performed with a read setup of paired end with a depth of 150 bp. Afterwards, raw data in FASTQ format were adapter- and quality trimmed. Mapping was performed against the Rattus norvegicus (mRatBN7.2.1110; January 05, 2024) genome sequence using Hisat2 alignment method. The mapped reads of each sample were assembled by StringTie (v1.3.3b) in a reference-based approach [22]. Differential expression analysis of two conditions/groups (three/four biological replicates per condition) was performed using the DESeq2 R package (1.42.0, [23]). Resulting p-values were corrected for multiple testing by false discovery rate (FDR) method and differentially expressed genes (DEGs) were filtered setting a threshold at the FDR adjusted p-value of 0.05 and a fold-change of ±1.5. The gene ontology (GO) analysis of differentially up- and downregulated genes was performed using Metascape platform using default parameters (R. norvegicus; February 2, 2024) or DAVID (R. norvegicus, February 2, 2024). Venn diagrams were generated using ggVenn R package (0.1.10).

### 2.3. Gene Set Enrichment Analysis

For Gene Set Enrichment Analysis (GSEA), the following published datasets were used. From [24], additional file 6 was downloaded and genes with an abs(logFC) > 0.15 and p_val_adj ≤ 0.05 of Mm3 (activated phagocytic microglia), Mm4 (Interferon response microglia), and Mm6 (Proliferative microglia) were extracted. From [25], supplementary Table S1 was downloaded and cluster 4 (Axon-tract associated microglia), cluster 7b (homeostatic microglia), cluster 8 (Aged microglia) and cluster 9 (Damage associated microglia) were extracted and used. From [3], supplementary Tables were downloaded and the first 100 genes for the immune cell cluster 0 (homeostatic microglia in MS), cluster 1 (microglial inflamed in MS – phagocytic) and cluster 8 (microglia inflamed in MS – immune response) were used. Related to astrocytes from [3], supplementary Tables were used and the first 100 genes for the astrocyte cluster cluster 3 (non-reactive astroglia in MS), cluster 1 (reactive astroglia in MS) and cluster 6 (inflamed astroglia in MS) were isolated. From [26], supplementary Table 3 was downloaded and genes with an abs(logFC) > 0.15 and p_val_adj ≤ 0.05 of cluster 5 (Astroglial signature upon EAE) were used for GSEA analysis. From [27], supplementary Table S6 was downloaded and genes with an abs(logFC) > 0.15 and p_val_adj ≤ 0.05 of cluster 8 (neuroinflammatory astrocytes) were isolated. After datasets were collected, gene lists were imported into the R environment and human and mouse gene names were translated to rodent gene nomenclature using Babelgene R package (22.9). Afterwards GSEA was performed using cluster profiler R package (4.10.0).

### 2.4. Statistics

Data are presented as mean values out of 3 (microglia) and 4 (astrocytes) biological replicates (n). Significance was accessed using the false discovery rate (FDR) method according to Benjamin and Hochberg and p-values/adjusted p-values are presented in this study as stated in the Figure legends. Graphical illustration with exception of Venn diagrams was performed using GraphPad Prism (9.3.0).

## 3. Results

### 3.1. HERV-W ENV drives microglial transcriptome to an immunologically activated state

In our previous studies, HERV-W ENV protein was shown to affect different glial cell types, driving them towards inflammatory and neurodegenerative phenotypes [14, 21]. To gain insights into the transcriptional signature of microglial cells upon HERV-W ENV exposure, primary microglia cells were stimulated with recombinant HERV-W ENV protein for 24h. Bulk mRNA sequencing revealed a total of 3397 significantly differentially expressed genes (DEGs) of which 1741 were downregulated and 1656 were upregulated (Fig. 1B). Interestingly, among the most significantly upregulated genes we detected lipocalin2 (Lcn2), a marker that was previously shown to be highly expressed by astroglial cells in the transgenic HERV-W ENV mouse model [21], as well as Nos2, the inducible nitric oxide-synthase that we previously reported to be a main mediator of neural toxicity (Fig. 1A)[14]. While upregulated DEGs where highly associated with proteins localized in membranes (Fig. 1C), downregulated genes were more related to the nucleus (Fig. 1D). Gene ontology (GO) biological process analyses supported this notion, as upregulated genes were mainly associated with immune response processes (Fig. 1E) whereas downregulated genes clustered primarily with decreased proliferation and DNA replication (Fig. 1F). Interestingly, upon HERV-W ENV stimulation downregulated genes were associated with gliogenesis (Fig. 1F). As this is defined as process that results in the generation of glial cells including the production of glial progenitors and their differentiation into mature glia, this observation indicates that ENV triggered microglial cells reduce their potential role in supporting glial cell development. KEGG pathway analysis of upregulated DEGs revealed that microglial genes were associated with three major pathways: Epstein-Bar virus infection, NOD-like receptor pathway, and Influenza A (Fig. 1G). All of these annotations involve different mechanisms and receptors with the potential of also being involved in HERV-W mediated activation processes. When it comes to the KEGG pathways that cluster with the downregulated genes, an association with cell cycle and replication pathways was observed (Fig. 1H), underlining the results of the biological pathway (GO) analysis.

**Figure 1:**
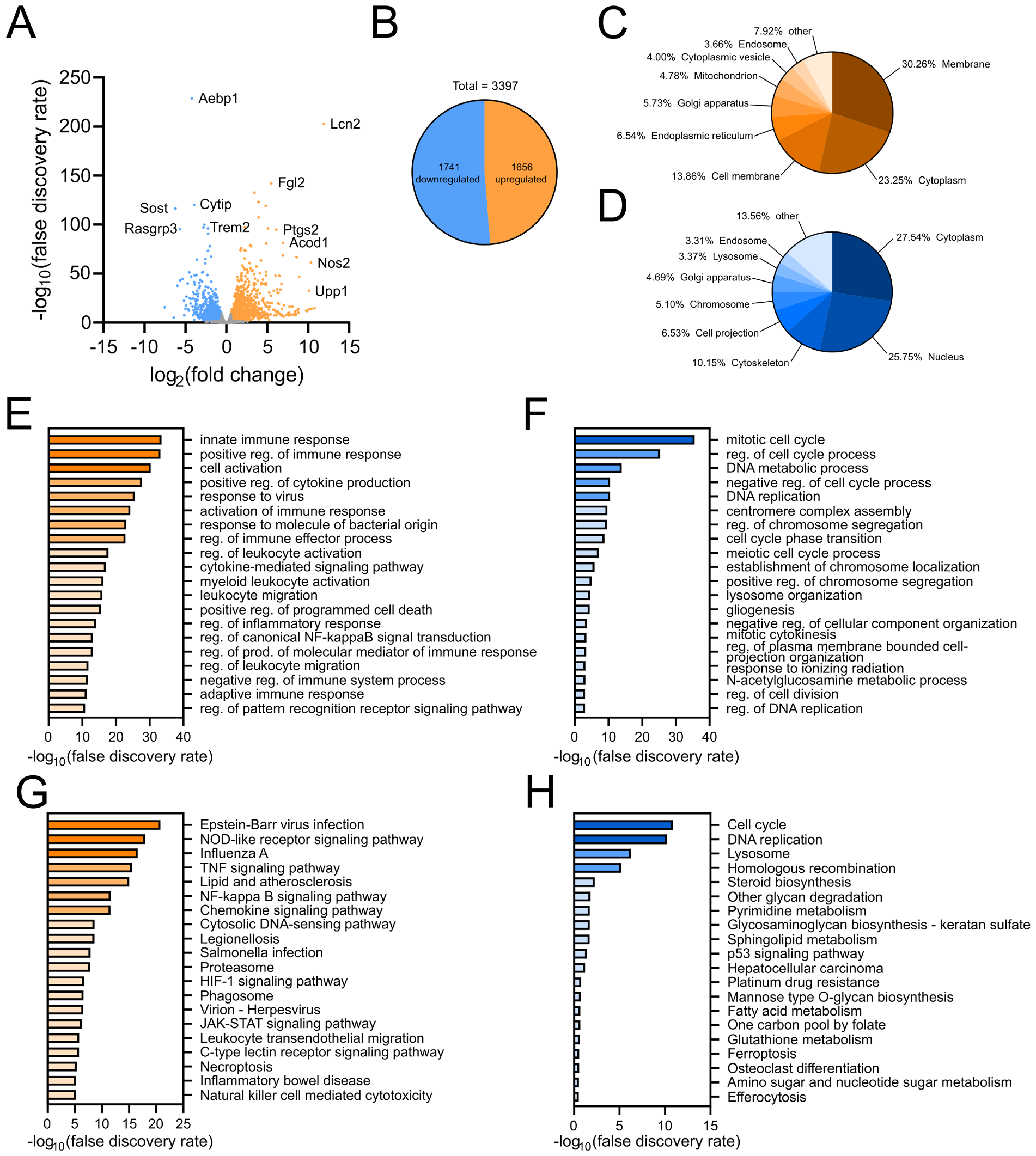
Transcriptome analysis of microglial cells stimulated with HERV-W ENV protein. (A) Vulcano plot showing log_2_(fold change) against -log_10_(false discovery rate) for the comparison of HERV-W ENV protein stimulated versus control-stimulated microglia. (B) Identification of 3397 DEGs (fold change of ± 1.5 and an FDR adjusted p-value of ≤ 0.05), of which 1741 were up- and 1656 were downregulated. (C, D) Cellular component analysis identifying the spatial association of up- (C) and downregulated genes (D). (E, F) Gene ontology analysis of Biological Processes identifying clusters of up- (E) and downregulated (F) genes. (G, H) KEGG pathway analysis revealing clusters of up- (G) and downregulated genes (H).

### 3.2. Gene expression differences between HERV-W ENV- and LPS stimulated microglial cells

HERV-W ENV was proposed to exert its effects mainly via TLR-4 signaling – shown at least for oligodendroglia, endothelia and immune cells [17, 20, 28]. Besides ENV, lipopolysaccharide (LPS) is known as one of the classical TLR4 activators. To reveal how specific and unique HERV-W ENV protein-evoked transcriptomic changes are, microglial cells were stimulated with 100 ng/ml LPS in parallel and equally subjected to bulk RNA sequencing. The analysis revealed a total number of 3760 differentially regulated genes (Fig. 2B), with Lcn2 and Nos2 still within the top upregulated genes (Fig. 2A). However, when comparing the LPS to HERV-W ENV evoked DEGs using a Venn diagram, substantial differences could be detected (Fig. 2C, supplementary Tables S1, S2). While 1261 upregulated genes and 1179 downregulated genes were detected in both groups, 395 genes were specifically upregulated in response to HERV-W ENV and 581 genes upon LPS treatment. Vice versa, we were able to identify 560 genes that were downregulated only upon HERV-W ENV exposure and 737 downregulated genes were detected exclusively when cells were stimulated with LPS. Interestingly, two genes (Bcar3, Sgk1) appeared to be upregulated upon LPS stimulation and downregulated upon HERV-W ENV exposure (Fig. 2C). To characterize the differences that are associated with these transcriptional disparities, gene ontology analysis for biological processes was performed using either the HERV-W-specific or the LPS-specific up- and downregulated genes as displayed by the Venn diagram. This disclosed that upregulated HERV-W-specific signatures were highly associated with adaptive immune system activation and lymphocyte recruiting (Fig. 2D), as opposed to LPS driven signatures - an interesting notion, indeed, as MS relapses are classically seen as lymphocyte driven. Among downregulated processes, again the term gliogenesis was also associated with the HERV-W ENV-specific response (Fig. 2E, compared to Fig. 1F). Furthermore, neurogenesis associated processes (regulation of neuron projection development; neuron projection development; regulation of synapse structure or activity) also clustered within the downregulated genes of the HERV-W ENV signature. After performing KEGG pathway analysis, additional differences between LPS- and ENV-stimulated cells were observed. This is exemplified by the Herpes simplex virus (HSV)-1 infection pathway emerging within HERV-W ENV upregulated genes and with LPS downregulated genes, indicating that additional receptors and their activation are involved in HERV-W ENV mediated signaling (Fig. 2F-G).

**Figure 2:**
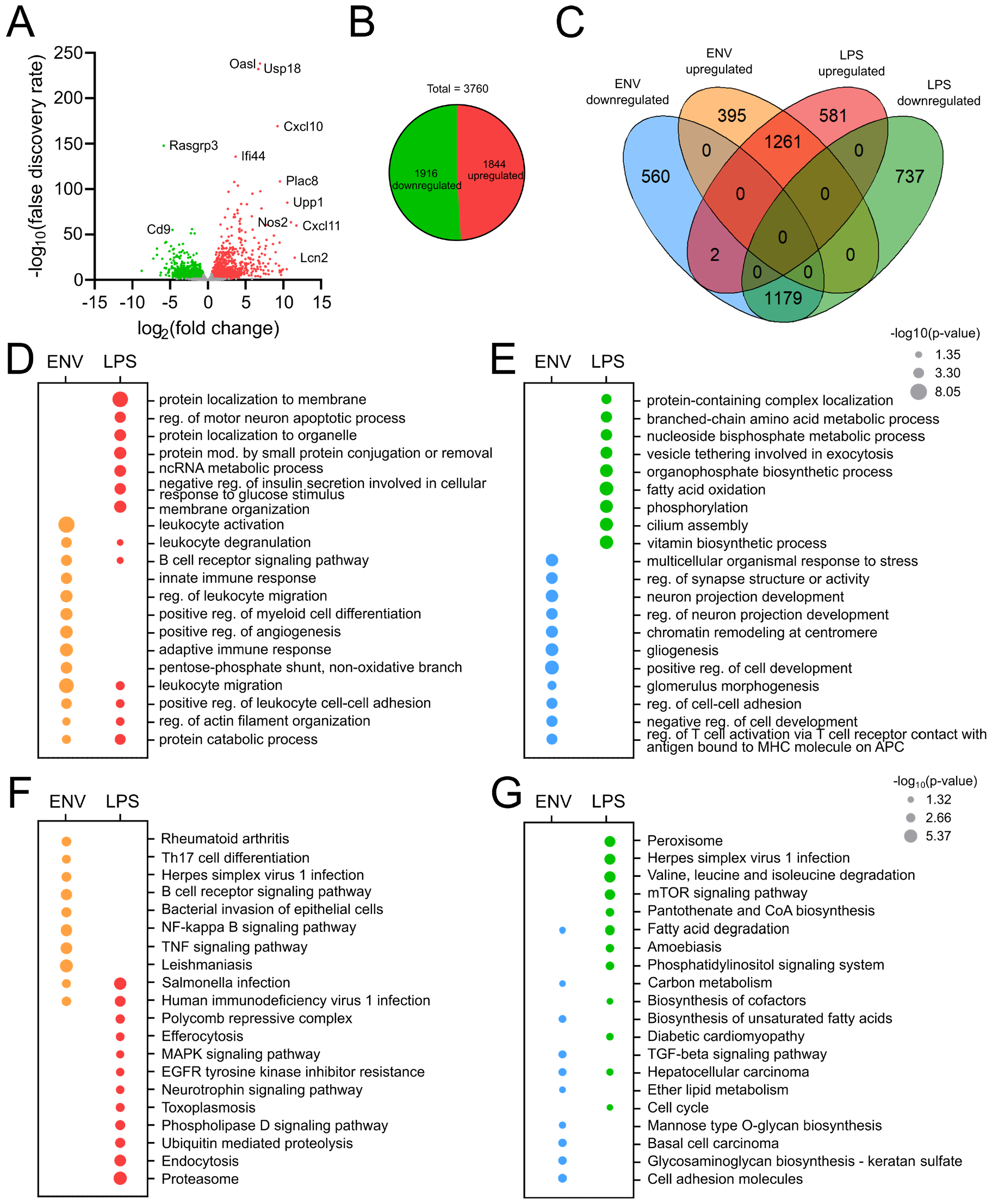
Transcriptional comparison of HERV-W ENV protein versus LPS stimulated microglia. (A) Vulcano plot showing log_2_(fold change) against -log_10_(false discovery rate) for the comparison of LPS protein stimulated versus control-stimulated microglia. (B) Identification of 3760 DEGs (fold change of ± 1.5 and an FDR adjusted p-value of ≤ 0.05), of which 1916 were up- and 1844 were downregulated. (C) Venn diagram of the identified DEGs from HERV-W ENV and LPS stimulated microglial cells. (D, E) Gene ontology analysis of Biological Processes identifying clusters of up- (D) and downregulated (E) genes upon exposure to HERV-W ENV versus LPS. (F, G) KEGG pathway analysis revealing clusters of up- (F) and downregulated genes (G) upon exposure to HERV-W ENV versus LPS. Dot sizes defines –log_10_(p-value).

### 3.3. Transcriptome analysis of astroglial cells upon HERV-W ENV stimulation

Besides previously presented reactions of microglia, oligodendroglial precursor cells, monocytes, lymphocytes and endothelial cells [14, 17, 19, 20] our recent *in vivo* analysis clearly revealed that also astrocytes directly react to HERV-W ENV exposure [21] indicated by the induction of neurotoxic marker expression and morphological transitions. To gain deeper insights into underlying transcriptional alterations of astroglia, bulk mRNA sequencing was performed after 24 hours of HERV-W ENV stimulation. Transcriptome analysis revealed a total of 2242 DEGs of which 1275 genes were downregulated and 967 genes were upregulated (Fig. 3B). Compared to the transcriptome response of microglial cells, it is therefore tempting to speculate that the astroglial response is milder since the total DEG number was lower (3397 in microglia, n=3 versus 2242 in astrocytes, n=4), while sequencing depth was similar. Again, Nos2 and Lcn2 were among the most significantly upregulated genes but also certain metalloproteases (Mmp9, Mmp3; Fig. 3A) were robustly induced. While upregulated genes of the microglial transcriptome were primarily associated with membranes, astrocytic upregulated genes were mostly associated with the cytoplasm (Fig. 3C). Interestingly, in contrast to microglia, upregulated genes of the astroglial transcriptome revealed an association with secreted proteins, indicating that astrocytes might play a key role in secreting trophic and/or inflammatory molecules – hence contributing to the generation of a neurotoxic environment as suggested recently [21]. On the other side, downregulated genes were mainly associated with cell membrane processes (Fig. 3D). After performing gene ontology clustering, upregulated astrocytic genes showed associations with innate immune responses (Fig. 3E) whereas, downregulated genes were associated with glial cell differentiation, neuron projection development and nervous system development (Fig. 3F), reflecting a decrease in their regenerative function while being polarized towards inflammation and neurodegeneration. Interestingly, both up- and downregulated genes showed associations with cell-cell adhesion. While downregulated genes were additionally highly associated with cell junction organization and extracellular matrix organization, it is tempting to speculate that certain cell-cell interactions might be decreased, whereas other cellular interactions might be increased. KEGG pathway analysis of upregulated DEGs revealed an association with cytokine-cytokine receptor interaction, primarily TNF signaling, as well as TLR signaling and viral infections such as EBV and Influenza A (Fig. 3G). When focusing on downregulated genes, KEGG pathway clustering revealed an association with cell adhesion molecules, axon guidance, calcium signaling and Wnt signaling.

**Figure 3:**
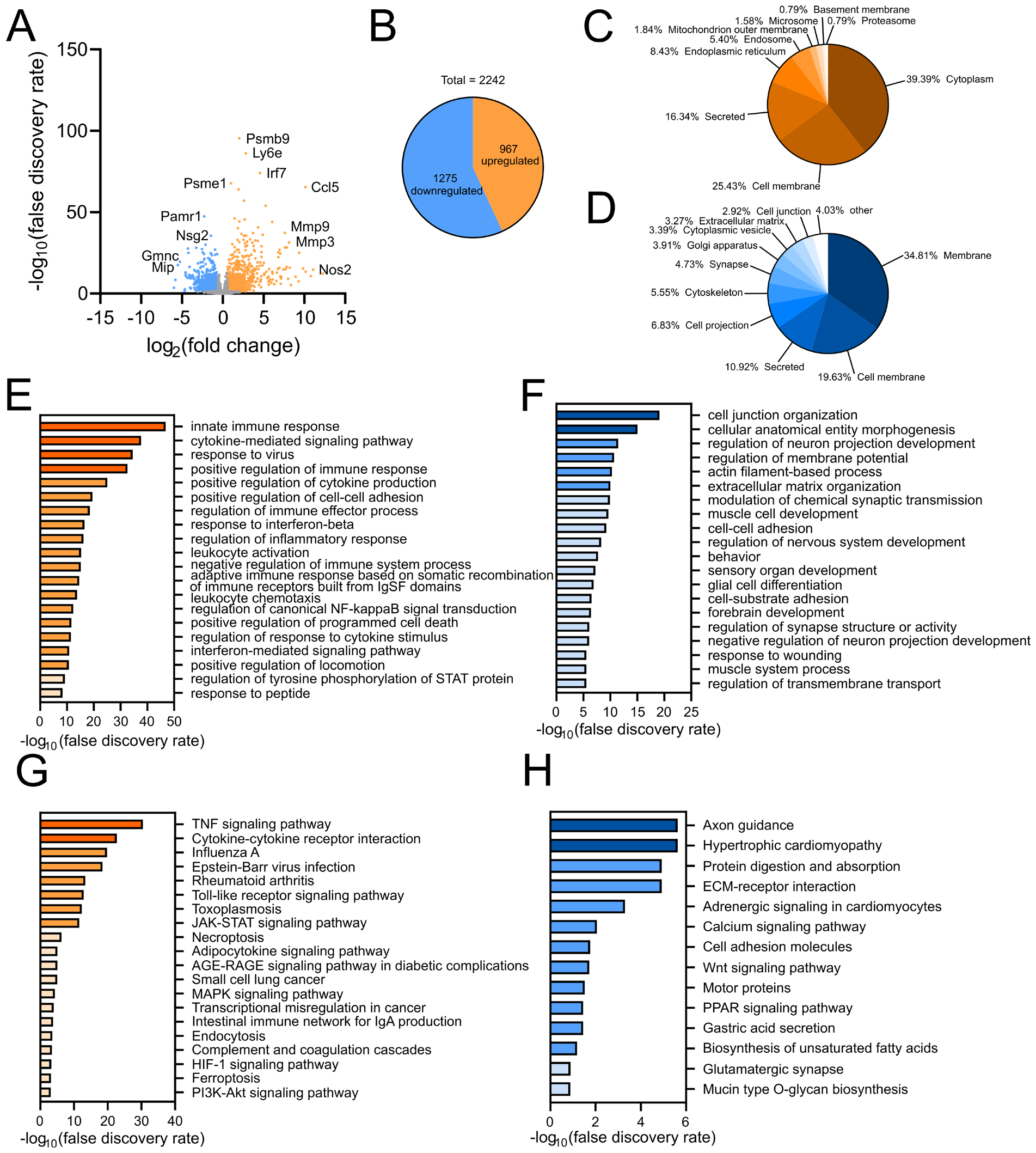
Transcriptome analysis of astroglial cells exposed to HERV-W ENV protein. (A) Vulcano plot showing log_2_(fold change) against -log_10_(false discovery rate) for the comparison of HERV-W ENV protein stimulated versus control-stimulated astrocytes. (B) Identification of 2242 DEGs (fold change of ± 1.5 and an FDR adjusted p-value of ≤ 0.05), of which 1275 were up- and 967 were downregulated. (C, D) Cellular component analysis identifying the spatial association of up- (C) and downregulated genes (D). (E, F) Gene ontology analysis of Biological Processes identifying clusters of up- (E) and downregulated (F) genes. (G, H) KEGG pathway analysis revealing clusters of up- (G) and downregulated genes (H).

### 3.4. Gene expression differences between HERV-W ENV- and LPS triggered astroglial cells

Similar to the determination of specific microglial transcriptome responses, we were also interested to identify differences between classical TLR4 activation by LPS and the activation mediated by HERV-W ENV. To this end, astroglial cells were stimulated with LPS and subsequent bulk mRNA sequencing was performed. The transcriptome analysis revealed a total number of 1349 differentially regulated genes of which 762 genes were downregulated and 587 genes were upregulated (Fig. 4B). Once more, Nos2 and Mmp9 could be detected among the most upregulated genes along with certain proinflammatory cytokines/chemokines such as (Il1b, Il1a, Ccl5, Cxcl11; Fig. 4A). As a next step, a Venn diagram was generated to identify similarities and differences between both astroglial expression patterns. Here, 535 genes were upregulated and 666 genes were downregulated in both groups, while 432 genes were specifically upregulated in response to HERV-W ENV as opposed to only 52 genes in the LPS paradigm (Fig. 4C; supplementary Tables S3, S4). Likewise, 609 genes were identified that were specifically downregulated upon HERV-W ENV exposure and only 96 genes were downregulated when cells were stimulated with LPS. Gene ontology analysis then disclosed that HERV-W ENV-specific gene inductions clustered primarily with pathways associated with the adaptive immune response and the attraction, recruiting and differentiation of lymphocytes (Fig. 4D). Upregulated genes of the LPS stimulated group showed no significant association within the top 20 biological processes. When analyzing both LPS- and HERV-W ENV downregulated genes, they clustered with cell-cell adhesion and cell junction behavior, indicating that these were not HERV-W-specific process (Fig. 4E). However, HERV-W ENV-specific downregulated genes were again associated with gliogenesis. KEGG pathway analysis revealed that upregulated astroglial genes specific to the exposure to HERV-W ENV protein were associated with complement and coagulation cascades (confirming our data on C3d induction *in vivo* [21]) as well as with cytokine-cytokine receptor interaction. HERV-W ENV-specific downregulated genes reflected axon guidance and motor protein involving process (Fig. 4F), indicating a reduced contribution to successful axonal regeneration.

**Figure 4:**
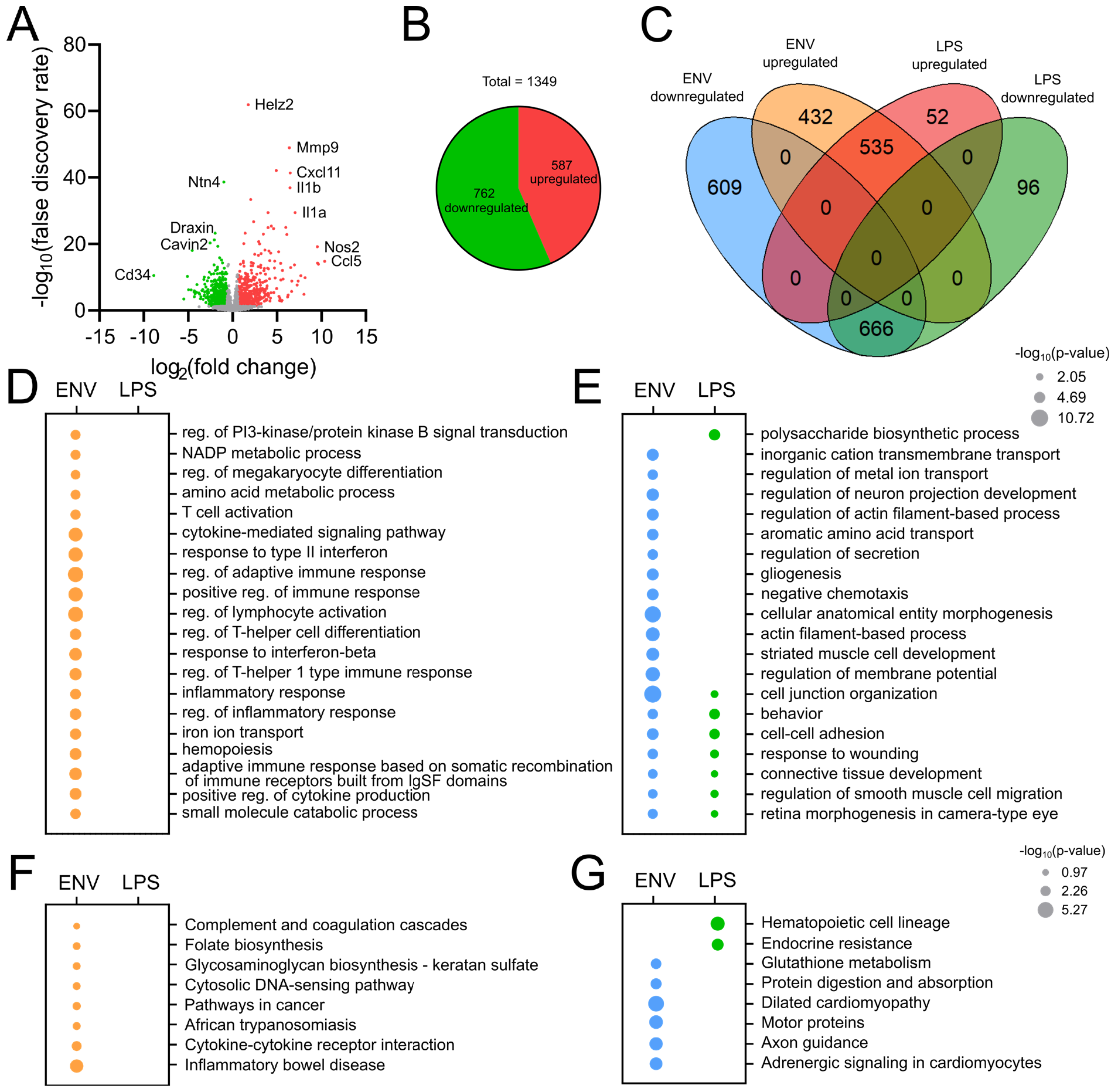
Comparison of HERV-W ENV-versus LPS mediated transcriptome changes of astroglial cells. (A) Vulcano plot showing log_2_(fold change) against -log_10_(false discovery rate) for the comparison of LPS stimulated versus control-stimulated astrocytes. (B) Identification of 1349 DEGs (fold change of ± 1.5 and an FDR adjusted p-value of ≤ 0.05), of which 762 were up- and 587 were downregulated. (C) Venn diagram of the identified DEGs from HERV-W ENV- and LPS stimulated microglial cells. (D, E) Gene ontology analysis of Biological Processes identifying clusters of up- (D) and downregulated (E) genes upon exposure to HERV-W ENV versus LPS. (F, G) KEGG pathway analysis revealing clusters of up- (F) and downregulated genes (G) upon exposure to HERV-W ENV versus LPS. Dot sizes defines –log_10_(p-value).

### 3.5. HERV-W dependent microglial signatures are enriched in MS

We next performed gene set enrichment analysis (GSEA) to compare the above presented genetic signatures with previously published gene patterns of disease associated microglia and astrocytes. In general, the comparison of HERV-W ENV dependent versus LPS mediated microglial transcriptomes with published datasets revealed that both groups showed an increased enrichment with disease and/or reactive signatures and a decreased enrichment with homeostatic microglial signatures (Fig. 5A). Here especially the “interferon responsive microglial” signature [24] evident by an HERV-W ENV related increased expression of Irf7, Usp18, Isg15 and Mx1 genes is currently of particular interest for the scientific community [29, 30]. On the other hand, homeostasis-associated genes such as P2RY12, Cx3cr1, Tmem119 and Tgfbr1 were downregulated [3, 24]. However, certain signatures also showed interesting differences when comparing HERV-W ENV vs. LPS stimulated microglia. This includes “axon-tract associated microglia” published by Hammond and colleagues [25] as well as “microglia inflamed in MS – immune responsive”, published by Absinta and colleagues [3], both of which were shown to be enriched in ENV stimulated microglia but decreased upon LPS exposure. While differences in “axon tract associated microglia” are mainly characterized by a change in Acr5, Adora3 and Csf1r expression levels, “immune responsive microglia inflamed in MS” were characterized by the induction of Cd74, Cd14 as well as C1qa, C1qb and C1qc genes. Interestingly, some of these markers have also been previously identified upon HERV-W ENV-signaling *ex vivo* and *in vivo* [6, 14].

**Figure 5:**
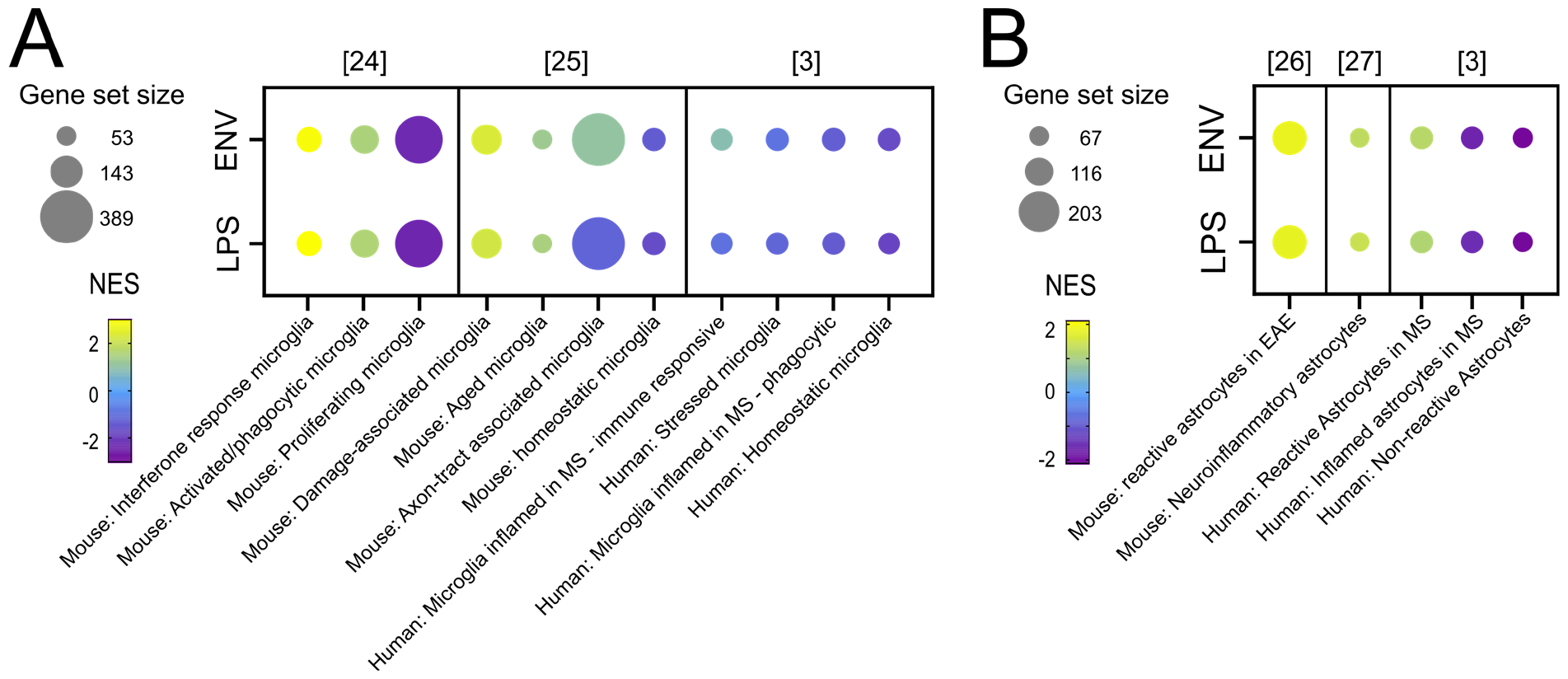
Gene set enrichment analysis of HERV-W ENV- vs. LPS stimulated microglial- and astroglial cells. (A) Gene set enrichment analysis (GSEA) of microglial transcriptomes upon HERV-W ENV or LPS stimulation. (B) Gene set enrichment analysis (GSEA) of astroglial transcriptomes upon HERV-W ENV or LPS stimulation. Dot sizes define the number of genes within the corresponding gene set and the colors relate to normalized enrichment score (NES). Numbers in brackets correspond to cited references.

Although fewer and less detailed publicly available astrocyte signature’s currently exist, a similar approach was used to compare the here disclosed astroglial transcriptomes to published datasets. The corresponding analysis disclosed that HERV-W ENV exposed astroglial cells are strongly associated with reactive [3, 26]and neuroinflammatory [27] signatures (related to induced Lcn2 Gbp2, C4b and Serping1 transcripts) whereas homeostatic signatures [3] were reduced (exemplified by Slc1a3, Aldh1a1, MerTK and Ptn gene downregulation). Of note, in contrast to the microglial transcriptomes, LPS stimulated astroglia revealed highly similar enrichment scores, indicating that astroglia react to HERV-W ENV in a less specific way and manner. A fact that might be related to a reduced receptor repertoire in astroglial cells as compared to microglia.

## 4. Discussion

The activation and expression of the HERV-W ENV protein has so far only been described in a few pathologies such as in MS, chronic inflammatory demyelinating polyradiculoneuropathy (CIDP), type-1 diabetes (T1D) as well as in some neurodevelopmental disorders [6, 31, 32]. Interestingly, recent evidence also strongly suggests its implication in COVID19 patients, further underlining the possible importance and impact of this endogenous viral entity [33]. It is therefore imperative to understand cellular responses towards HERV-W ENV in greater detail to advance the development of future therapeutic approaches limiting its pathological impact.

While we were recently able to characterize important tissue de- and regeneration effects of HERV-W ENV in a novel *in vivo* model [21], thereby recapitulating and confirming functions that were previously suggested from primary cell culture models or by clinical observations on patients treated with a HERV-W ENV directed neutralizing antibody termed Temelimab [34], a different approach was conducted here to understand its mode of action and to determine how specific and unique this pathological entity’s signaling is. We were able to describe for the first-time complete transcriptomes of HERV-W ENV-exposed microglial- and astroglial cells, identifying many inflammation-associated genes that are strongly upregulated (Fig. 1,3). Furthermore, by comparing these HERV-W ENV-dependent signatures with classical LPS-mediated TLR4 signaling activation, important differences were recognized that underline a specific mode of action (Fig. 2,4). In this regard, it was found that HERV-W ENV leads to an induction of adaptive immune responses and to a decrease in gliogenesis- as well as neuroregeneration-associated genes in both cell populations - notably in contrast to the LPS mediated activation (Fig. 2D,E and Fig. 4D,E). This is an important notion since all of these pathways play important roles in the pathology of MS. In addition, astrocytes revealed to respond by impaired cell-adhesion-, cell/cell-junction- as well as Wnt signaling parameters (Fig. 3). In this regard the observed upregulation of cell-adhesion molecule encoding genes such as Icam1, Vcam1 as well as of integrins Itgal, Itgb8, Itgax, Itgb2 and cadherins Cdh4, Cdh13 along downregulation of Ncam1, Cdh20, Cdh8, Itga11, Itga1, Itga4, Itgb4 and Itgb5 transcripts is of particular interest. Since members of these genes families are involved in maintaining the blood-brain-barrier (BBB) integrity [35], it is tempting to speculate that its function is disturbed. Such a functional impact is corroborated by the observed negative influence of HERV-W ENV towards Wnt signaling (Fig. 3H). Wnt factor release from astrocytes was repeatedly shown to be crucial for BBB maintenance [36, 37] and Wnt7a, which is seen as the main driver for BBB integrity [38], was significantly downregulated in response to HERV-W ENV. As an activating and inflammatory effect of the ENV protein exerted onto endothelial cells has previously been described [17], this suggests on overall negative effect on BBB integrity and function.

Besides the observed differences between microglial- and astroglial responses, we also noticed that HERV-W and LPS transcriptomes were also different, however, in a cell-specific way. Microglia appear to react more extensively towards ENV protein exposure as compared to astroglia and this reaction also differed significantly more from LPS mediated signatures (Fig. 2C compared to Fig. 4C), indicating that microglial signaling might involve multiple pathways and receptors. Apart from TLR4, retroviral envelope protein interactions were shown for TLR2 [39], MFSD2 [40], ASCT1/Slc1a4 [41], ASCT2/Slc1a5 [42] and MCT-1/Slc16a1 [43]. Our data so far only confirm the expression of TLR4 on microglial cells [14] but to what degree such other receptors additionally contribute to the observed neurodegenerative phenotype acquisition needs to be addressed in future experiments. Nevertheless, a preliminary assessment of gene expression according to [44] confirmed microglial CD14, TLR2 and TLR4 expression but also revealed strong FPKM values for ASCT2/Slc1a5 and trace FPKM values of ASCT1/Slc1a4 and MCT-1/Slc16a1 (data not shown).

Recent transcriptome studies accurately described the diversity of microglial- and astroglial subphenotypes in different diseases and mouse models [3, 24, 25]. We therefore compared the here generated transcriptomes to existing microglial- and astroglial signatures (Fig. 5) aiming at a description of HERV-W’s specific mode of action in health and disease. Compared to LPS-mediated effects, HERV-W ENV stimulated microglial signatures were characterized by an enrichment in “axon tract associated microglia” [25]. This molecular assignment fits well to our previous documentation that in chronic active MS lesions as well as in a functional *ex vivo* model, HERV-W ENV triggered microglia were closely associated with myelinated axons. Moreover, secreted protein analysis in the applied *ex vivo* model then confirmed axon damage- and myelin proteins to be released in response to these activated microglial cells [14]. Also, a HERV-W ENV-specific association with the “immune responsive microglial inflamed in MS” [3] gene cluster was identified. As this dataset directly derives from MS patients, it demonstrates an important difference between these two pathological triggers and strongly supports the notion that HERV-W is an MS-specific pathological entity. Interestingly, within this cluster the main differentially expressed genes were Cd74, Cd14 as well as C1qa, C1qb and C1qc, most of them had recently been reported to be induced in the transgenic mouse model [21], thus cross-validating these effects. Finally, the fact that both, LPS and HERV-W ENV mediated transcriptomes, correlated with the “interferon responsive microglial”[24] signature is also of interest, since this particular cellular phenotype is currently discussed as one of the main mediators of neurodegeneration [29, 30] along with its already accepted association with viral infections and recruitment of adaptive immune cells (summarized by [45]).

While one limitation of this study relates to the fact that the experiments had to be performed *ex vivo* in order to decipher HERV-W from LPS mediated effects, the here outlined comparisons to datasets derived from patients and/or *in vivo* models identified large overlaps hence supporting the validity of the chosen approach. Upcoming studies using differentially addressed gene clusters on human (MS) tissue samples and/or the recently developed transgenic mouse model will be used to further corroborate our new findings. Likewise, it is worth mentioning that all so far published astroglial signatures do not decipher a possible influence or modulation by microglia and lymphocytes although such cell/cell interactions have already been described [3, 46]. Even though, the here presented evidence suggests that HERV-W ENV’s functionality in astrocytes is restricted to TLR4 activation only, more specific consequences mediated via microglial- or lymphocyte crosstalk could be expected.

Overall, the here presented data strongly supports the notion that the concomitant activation of both glial cell types by the ENV protein leads to the generation of a neurotoxic environment. Such a milieu is characterized by inhibited regenerative processes and by enhanced neurodegeneration as recently observed in the transgenic HERV-W ENV expressing mouse model [6] and as deduced from analysis of Temelimab treated patients [34].

## 5. Conclusion

We can conclude that cellular reactions in response to the HERV-W ENV protein are indeed specific and unique, supporting its characterization as an important pathological factor that deserves more attention. Taking this ENV protein’s presence in several neurodegenerative and developmental diseases into account, it is tempting to speculate that it even constitutes a common factor related to chronic inflammation and neurodegeneration – including the smoldering neuroinflammation process as it is currently in focus of MS research [47].

## 6. Author contributions

JG, LW and LR performed *in vitro* experiments

JG, FH and JS performed RNA sequencing analysis

JG, LR, UM, PK were involved in data interpretation

JG and PK wrote the manuscript

LR and UM reviewed the manuscript

## 7. Funding

Research presented in this manuscript was financially supported by the Christiane and Claudia Hempel Foundation for regenerative medicine and by the James and Elisabeth Cloppenburg, Peek and Cloppenburg Düsseldorf Stiftung.

## 8. Data availability Statement

RNASeq datasets generated for this study are publicly available in the NIH Gene Expression Omnibus (GEO) repository: https://www.ncbi.nlm.nih.gov/geo/, accession number.

## 9. Acknowledgement

We acknowledge the support of Brigida Ziegler and Birgit Blomenkamp-Radermacher for their technical support. Furthermore, we would like to thank Geneuro SA for providing recombinant HERV-W ENV protein.

## 10. Conflict of interest

All authors declare that the research was conducted in the absence of any commercial or financial relationships that could be construed as a potential conflict of interest.

## Notes

### Competing Interest Statement

The authors have declared no competing interest.

## References

1. Yong, H.Y.F. and V.W. Yong, Mechanism-based criteria to improve therapeutic outcomes in progressive multiple sclerosis. Nat Rev Neurol, 2022. 18(1): p. 40–55.

2. Reeves, J.A., et al., Reliability of paramagnetic rim lesion classification on quantitative susceptibility mapping (QSM) in people with multiple sclerosis: Single-site experience and systematic review. Mult Scler Relat Disord, 2023. 79: p. 104968.

3. Absinta, M., et al., A lymphocyte-microglia-astrocyte axis in chronic active multiple sclerosis. Nature, 2021. 597(7878): p. 709–714.

4. Escartin, C., et al., Reactive astrocyte nomenclature, definitions, and future directions. Nat Neurosci, 2021. 24(3): p. 312–325.

5. Paolicelli, R.C., et al., Microglia states and nomenclature: A field at its crossroads. Neuron, 2022. 110(21): p. 3458–3483.

6. Gruchot, J., et al., Interplay between activation of endogenous retroviruses and inflammation as common pathogenic mechanism in neurological and psychiatric disorders. Brain Behav Immun, 2023. 107: p. 242–252.

7. Perron, H., et al., Leptomeningeal cell line from multiple sclerosis with reverse transcriptase activity and viral particles. Res Virol, 1989. 140(6): p. 551–61.

8. Mameli, G., et al., Activation of MSRV-type endogenous retroviruses during infectious mononucleosis and Epstein-Barr virus latency: the missing link with multiple sclerosis? PLoS One, 2013. 8(11): p. e78474.

9. Mameli, G., et al., Expression and activation by Epstein Barr virus of human endogenous retroviruses-W in blood cells and astrocytes: inference for multiple sclerosis. PLoS One, 2012. 7(9): p. e44991.

10. Bjornevik, K., et al., Longitudinal analysis reveals high prevalence of Epstein-Barr virus associated with multiple sclerosis. Science, 2022. 375(6578): p. 296–301.

11. Garson, J.A., et al., Detection of virion-associated MSRV-RNA in serum of patients with multiple sclerosis. Lancet, 1998. 351(9095): p. 33.

12. Mameli, G., et al., Novel reliable real-time PCR for differential detection of MSRVenv and syncytin-1 in RNA and DNA from patients with multiple sclerosis. J Virol Methods, 2009. 161(1): p. 98–106.

13. Perron, H., et al., Human endogenous retrovirus type W envelope expression in blood and brain cells provides new insights into multiple sclerosis disease. Mult Scler, 2012. 18(12): p. 1721–36.

14. Kremer, D., et al., pHERV-W envelope protein fuels microglial cell-dependent damage of myelinated axons in multiple sclerosis. Proc Natl Acad Sci U S A, 2019. 116(30): p. 15216–15225.

15. van Horssen, J., et al., Human endogenous retrovirus W in brain lesions: Rationale for targeted therapy in multiple sclerosis. Mult Scler Relat Disord, 2016. 8: p. 11–8.

16. Perron, H., et al., Human endogenous retrovirus protein activates innate immunity and promotes experimental allergic encephalomyelitis in mice. PLoS One, 2013. 8(12): p. e80128.

17. Duperray, A., et al., Inflammatory response of endothelial cells to a human endogenous retrovirus associated with multiple sclerosis is mediated by TLR4. Int Immunol, 2015. 27(11): p. 545–53.

18. Perron, H., et al., Multiple sclerosis retrovirus particles and recombinant envelope trigger an abnormal immune response in vitro, by inducing polyclonal Vbeta16 T-lymphocyte activation. Virology, 2001. 287(2): p. 321–32.

19. Rolland, A., et al., The envelope protein of a human endogenous retrovirus-W family activates innate immunity through CD14/TLR4 and promotes Th1-like responses. J Immunol, 2006. 176(12): p. 7636–44.

20. Kremer, D., et al., Human endogenous retrovirus type W envelope protein inhibits oligodendroglial precursor cell differentiation. Ann Neurol, 2013. 74(5): p. 721–32.

21. Gruchot, J., et al., Transgenic expression of the HERV-W envelope protein leads to polarized glial cell populations and a neurodegenerative environment. Proc Natl Acad Sci U S A, 2023. 120(38): p. e2308187120.

22. Pertea, M., et al., StringTie enables improved reconstruction of a transcriptome from RNA-seq reads. Nat Biotechnol, 2015. 33(3): p. 290–5.

23. Love, M.I., W. Huber, and S. Anders, Moderated estimation of fold change and dispersion for RNA-seq data with DESeq2. Genome Biol, 2014. 15(12): p. 550.

24. Miedema, A., et al., Brain macrophages acquire distinct transcriptomes in multiple sclerosis lesions and normal appearing white matter. Acta Neuropathol Commun, 2022. 10(1): p. 8.

25. Hammond, T.R., et al., Single-Cell RNA Sequencing of Microglia throughout the Mouse Lifespan and in the Injured Brain Reveals Complex Cell-State Changes. Immunity, 2019. 50(1): p. 253–271 e6.

26. Wheeler, M.A., et al., MAFG-driven astrocytes promote CNS inflammation. Nature, 2020. 578(7796): p. 593–599.

27. Hasel, P., et al., Neuroinflammatory astrocyte subtypes in the mouse brain. Nat Neurosci, 2021. 24(10): p. 1475–1487.

28. Madeira, A., et al., MSRV envelope protein is a potent, endogenous and pathogenic agonist of human toll-like receptor 4: Relevance of GNbAC1 in multiple sclerosis treatment. J Neuroimmunol, 2016. 291: p. 29–38.

29. Roy, E. and W. Cao, Glial interference: impact of type I interferon in neurodegenerative diseases. Mol Neurodegener, 2022. 17(1): p. 78.

30. Roy, E.R., et al., Type I interferon response drives neuroinflammation and synapse loss in Alzheimer disease. J Clin Invest, 2020. 130(4): p. 1912–1930.

31. Küry, P., et al., Human Endogenous Retroviruses in Neurological Diseases. Trends Mol Med, 2018. 24(4): p. 379–394.

32. Levet, S., et al., An ancestral retroviral protein identified as a therapeutic target in type-1 diabetes. JCI Insight, 2017. 2(17).

33. Charvet, B., et al., SARS-CoV-2 awakens ancient retroviral genes and the expression of proinflammatory HERV-W envelope protein in COVID-19 patients. iScience, 2023. 26(5): p. 106604.

34. Hartung, H.P., et al., Efficacy and safety of temelimab in multiple sclerosis: Results of a randomized phase 2b and extension study. Mult Scler, 2022. 28(3): p. 429–440.

35. Kadry, H., B. Noorani, and L. Cucullo, A blood-brain barrier overview on structure, function, impairment, and biomarkers of integrity. Fluids Barriers CNS, 2020. 17(1): p. 69.

36. Daneman, R., et al., Wnt/beta-catenin signaling is required for CNS, but not non-CNS, angiogenesis. Proc Natl Acad Sci U S A, 2009. 106(2): p. 641–6.

37. Guerit, S., et al., Astrocyte-derived Wnt growth factors are required for endothelial blood-brain barrier maintenance. Prog Neurobiol, 2021. 199: p. 101937.

38. Zhou, Y., et al., Canonical WNT signaling components in vascular development and barrier formation. J Clin Invest, 2014. 124(9): p. 3825–46.

39. Reuven, E.M., et al., The HIV-1 envelope transmembrane domain binds TLR2 through a distinct dimerization motif and inhibits TLR2-mediated responses. PLoS Pathog, 2014. 10(8): p. e1004248.

40. Esnault, C., et al., A placenta-specific receptor for the fusogenic, endogenous retrovirus-derived, human syncytin-2. Proc Natl Acad Sci U S A, 2008. 105(45): p. 17532–7.

41. Antony, J.M., et al., The human endogenous retrovirus envelope glycoprotein, syncytin-1, regulates neuroinflammation and its receptor expression in multiple sclerosis: a role for endoplasmic reticulum chaperones in astrocytes. J Immunol, 2007. 179(2): p. 1210–24.

42. Blond, J.L., et al., An envelope glycoprotein of the human endogenous retrovirus HERV-W is expressed in the human placenta and fuses cells expressing the type D mammalian retrovirus receptor. J Virol, 2000. 74(7): p. 3321–9.

43. Blanco-Melo, D., R.J. Gifford, and P.D. Bieniasz, Co-option of an endogenous retrovirus envelope for host defense in hominid ancestors. Elife, 2017. 6.

44. Zhang, Y., et al., An RNA-sequencing transcriptome and splicing database of glia, neurons, and vascular cells of the cerebral cortex. J Neurosci, 2014. 34(36): p. 11929–47.

45. Mertowska, P., et al., Immunomodulatory Role of Interferons in Viral and Bacterial Infections. Int J Mol Sci, 2023. 24(12).

46. Liddelow, S.A., et al., Neurotoxic reactive astrocytes are induced by activated microglia. Nature, 2017. 541(7638): p. 481–487.

47. Giovannoni, G., et al., Smouldering multiple sclerosis: the ‘real MS’. Ther Adv Neurol Disord, 2022. 15: p. 17562864211066751.

